# Exploring differentially expressed key genes related to development of follicle by RNA-seq in Peking ducks (*Anas Platyrhynchos*)

**DOI:** 10.1101/483818

**Authors:** Jindong Ren, Changsen Sun, Li Chen, Jianhong Hu, Xuetao Huang, Xiaolin Liu, Lizhi Lu

## Abstract

Duck follicle enter different reproductive phases throughout life, and follicle gene expression patterns differ according to these phases. In particular, differentially expressed genes and related to development of follicle (mRNAs) play an important role to explore the key genes in this process; however, the expression profiles of these genes remain unclear. In this study, transcriptome sequencing was used to investigate the expression levels of duck ovarian genes, and comparative transcriptional analysis was carried out to identify differential genes, cluster them into groups and function identification. The results showed differential expression of 593 coding genes between young and laying ducks, and of 518 coding genes between laying and old ducks. In further GO analysis, 35 genes from the comparison bewtween old ducks and laying ducks have significant been changed involved in hormones related to follicle development. They include up-regulated genes StAR, CYP17, EPOX, 3β-HSD, CYP1B1 CYP19A1 and down-regulated genes SR-B1 in laying ducks hormone synthesis than old ducks. Among which EPOX is a key gene for time special highly expression during egg laying stage, and other key regulatory genes’ highly expression showed in young and laying stage and lower expression showing with follicular development stopping. Therefore, EPOX is key regulator for duck follicle development in laying period, when its expression level decrease 98% the follicular development will stopping in duck life cycle.

## INTRODUCTION

Maintenance of the physiological status of ovaries at different times requires serial specific genes and some biology molecular, such as regulation element function protein. For egg-laying poultry, the ovary follicle is characterized by three life phases: the growth phase, laying phase, and maternity phase. During the growth period, the primordial follicle is assembled and prepared for egg-laying at sexual maturation, and remains in a quiescent state [1]. During the egg-laying period, the follicle is activated and ovulation becomes the main activity of the ovary, regulated by the secretion of sex hormones [2]. In contrast to the egg-laying period, most reproductive activities cease during the maternity phase, including the secretion of sexual hormones. The presence of three contrasting ovary phases in poultry indicates that different genes associated with each phase are expressed differentially in clusters. The present study aimed to identify these differentially expressed gene clusters and their function to provide a molecular-level understanding of the different developmental phases evident in duck ovaries.

The follicle in ovary of poultry develop dynamically throughout life, beginning at gametogenesis. During the multiplicative phase, the assembly of the primordial follicles is completed [3]. Throughout this process, most primordial follicles are approximately 0.05 mm in diameter and remain in a quiescent state until sexual maturation [1, 4]; a special characteristic of this phase is a change in the shape of granulosa cells from flat to cuboidal. The gene promoting the transition from quiescence to slow growing follicles is yet to be investigated in poultry; however, the anti-Mullerian hormone (AMH) and KIT-ligand have been identified as potential factors in the analogous transition in mammals [5-7]. After the multiplicative phase, the ovary begins follicular development in the egg-laying period. During this period, the pulsatile secretion of gonadotropin-releasing hormone (GnRH) stimulates the pituitary gonadotropin secretion [1], and causing follicles to be selected to enter the follicle hierarchy. As in mammals, follicle stimulating hormone (FSH) is an important factor in the development of pre-hierarchical follicles in poultry [8], and it induces follicular selection [9]. Although the signal pathway of FSH is known, the precise role of FSH is unclear, especially in the transition from the egg-laying phase to the post-laying phase. Clock genes, such as *BMAL1-CLOCK* (Brain and Muscle ARNT-like-1), are the only genes involved in phase transitions investigated to date [10]. However, the mechanism underlying the regulation of transition by the clock genes is unclear [11-12], and which signal path play a key function in follicle development and change that also is unclear, in previously.

Therefore, in this study, we identified and quantified genes from duck follicle in three life stages, to identify the potential key genes involved in transitions between the stages by comparing differential gene expression profiles in the three phases, and by examining molecular markers that may be important for egg laying in ducks.

## MATERIALS AND METHODS

### Ethics statement

The protocol for the care and slaughter of experimental ducks was approved by the Institutional Animal Care and Use Committee of the Northwest Agriculture and Forestry University, and carried out in accordance with the Regulations for the Administration of Affairs Concerning Experimental Animals (China, 1988). All protocols were carried out according to recommendations proposed by the Animal welfare European Commission (1997), and all efforts were made to minimize the suffering of experimental ducks.

### Tissue sampling and RNA preparation

Ovary follicle samples were collected from the Zhongwang Beijing Duck Breeding Farm (Huzhou, China). Three birds each from three different age groups (60 day old young ducks, YD; 160 day old first-laying ducks, FL; and 490 day old stop-laying ducks, OD) were slaughtered for tissue sampling. Fresh ovary follicle samples were washed in phosphate-buffered saline (PBS; Gibco, Fisher Scientific, Waltham, MA, USA) and immediately frozen in liquid nitrogen. Total RNA was extracted from each sample using an Agilent 2100 RNA Nano 6000 Assay Kit (Agilent Technologies, Santa Clara, CA, USA). RNA concentration and purity was determined using a spectrophotometer to investigate the OD260/OD280 of sample (NanoVue; GE Healthcare, Piscataway, NJ, USA).

### Sequencing and assembly of expressed genes

A total quantity of 3 μg RNA per sample was used as initial material for RNA sample preparation. Ribosomal RNA was extracted using an Epicentre Ribo-Zero™ Gold Kit (Epicentre Technologies, Madison, WI, USA) and was used to generate sequencing libraries with varied index label using a NEBNextUltra™ Directional RNA Library Prep Kit for Illumina (New England BioLabs, Ipswich, MA, USA) according to the manufacturer’s instructions. The clustering of index-coded samples was carried out on a cBot cluster generation system using TruSeq PE Cluster Kit v3-cBot-HS (Illumina, San Diego, CA, USA) according to manufacturer’s instructions. After cluster generation, the libraries were sequenced on an Illumina Hiseq 4000 platform (Illumina, San Diego, CA, USA) and 150 bp paired-end reads were generated. Clean data were obtained by filtering the raw reads and removing polluted reads, low-quality reads, and reads with unknown bases, accounting for more than 5% of total genomic material. Filtered reads were aligned to the duck genome using TopHat version 2.0.12 [13]and mapped with Bowtie 2 version 2.2.3 [14].

### Quantification of gene expression levels and differential analysis

The number of clean reads per gene per sample was counted using HTSeq version 0.6.0 [15]. Reads per kilobase per million mapped reads (RPKM) were calculated to estimate the expression level of genes in each sample using the formula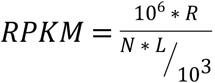, where *R* is the number of reads in a sample assigned to a gene, *N* is the total number of mapped reads in the sample, and *L* is the length of the gene [16].

Pairwise analyses of differential gene expression were carried out with biological replicates using DEGseq version 1.16. The gene expression levels were determined using a negative binomial distribution model, and genes with *q* ≤ 0.05 and |log2ratio| ≥ 1 were identified as differentially expressed genes (DEGs) [17]. The expression levels of all divergent genes were log2-transformed, the Euclidean distance was calculated, and clustering was performed using Hierarchical Cluster methods in R software (version 3.2.5).

### GO and KEGG enrichment analysis of DEGs

The GO (Gene Ontology, http://geneontology.org/) enrichment of DEGs was implemented by the hypergeometric test, in which p-value is calculated and adjusted as q-value, and data background is genes in the whole genome. GO terms with q<0.05 were considered to be significantly enriched [18].

KEGG (Kyoto Encyclopedia of Genes and Genomes, http://www.kegg.jp/) enrichment of DEGs was implemented by the hypergeometric test, in which p-value was adjusted by multiple comparisons as q-value. KEGG terms with q<0.05 were considered to be significantly enriched [19].

## RESULTS

### Differentially expressed genes

High-throughput sequencing technology was used to analyze the gene expression profiles of ovary follicle tissue from the three sampled life stages. Reads of each RNA sample from the three groups were identified and annotated based on available duck genomic data. This assembly identified 15576 genes. The reads for each gene were used to calculate its expression level, as indicated by RPKM. Based on the significance values, 593 and 518 differentially expressed genes were detected in the comparative analyses between FL and YD, and OD and FL, respectively.

### Divergent gene clusters

The expression levels of the divergent genes identified in pairwise comparisons were clustered in distinct groups. Among the divergent genes identified in the comparison between FL and YD, five genes were significantly downregulated with age and formed a group clustered by expression levels in young ducks (Cluster No. 1, Table 1; divergent gene names, descriptions, and cluster numbers are provided in the supplementary tables 1.); two genes showed significant swing expression patterns (i.e., high, low, and high expression levels over time), rather than consistent levels patterns and formed another cluster group (Cluster No. 2, Fig 1C.); and the remaining genes showed consistent expression levels and formed a third cluster group (Cluster No. 3). The divergent genes identified in the comparison between first-laying and old duck tissue showed a similar pattern, although Cluster No. 3 in laying ducks showed significant swing expression levels over time rather than remaining consistent. The genes from Cluster No. 2 in young ducks showed similar expression levels; however, a pattern of upregulation was identified in these genes (Fig 1A.). The most suitable clustering schema was derived from old duck expression data; however, the most suitable clustering schema was recovered by excluding the gene with the highest expression level in old ducks.

**Table 1.** Expression levels in young ducks (YD), first-laying ducks (FL) and old ducks (OD) for genes differentially expressed between FL and YD. “Ratio of SS” indicates the ratio of within-cluster sum of squares to total sum of squares. Different lowercase letters in the same row indicate significant difference at P < 0.05.

| Cluster source | Cluster No. | Number of divergent genes | Ratio of SS | YD | FL | OD |
| --- | --- | --- | --- | --- | --- | --- |
| Young ducks | 1 | 5 |  | 559.32<sup>a</sup> ± 263.946 | 119.65<sup>b</sup> ± 105.418 | 178.42<sup>b</sup> ± 184.657 |
|  | 2 | 586 | 84.4% | 12.65 ± 25.233 | 11.06 ± 20.629 | 11.46 ± 25.191 |
|  | 3 | 2 |  | 1979.20<sup>a</sup> ± 282.783 | 290.44<sup>a</sup> ± 325.376 | 927.7<sup>ab</sup> ± 1173.049 |
| Laying ducks | 1 | 3 |  | 1185.83 ± 919.616 | 326.48 ± 168.034 | 821.78 ± 818.713 |
|  | 2 | 53 | 62.4% | 95.23 ± 246.775 | 65.23 ± 31.433 | 59.6 ± 59.423 |
|  | 3 | 537 |  | 10.36<sup>a</sup> ± 33.695 | 6<sup>b</sup> ± 8.503 | 7.15<sup>a</sup> ± 11.844 |
| Old ducks | 1 | 1 |  | 2179.15 | 520.51 | 1757.17 |
|  | 2 | 9 | 93.6% | 226.29 ± 234.107 | 120.53 ± 59.735 | 214.33 ± 122.279 |
|  | 3 | 583 |  | 17.07 ± 83.622 | 10.38 ± 18.869 | 9.91 ± 16.162 |

**Fig 1.**
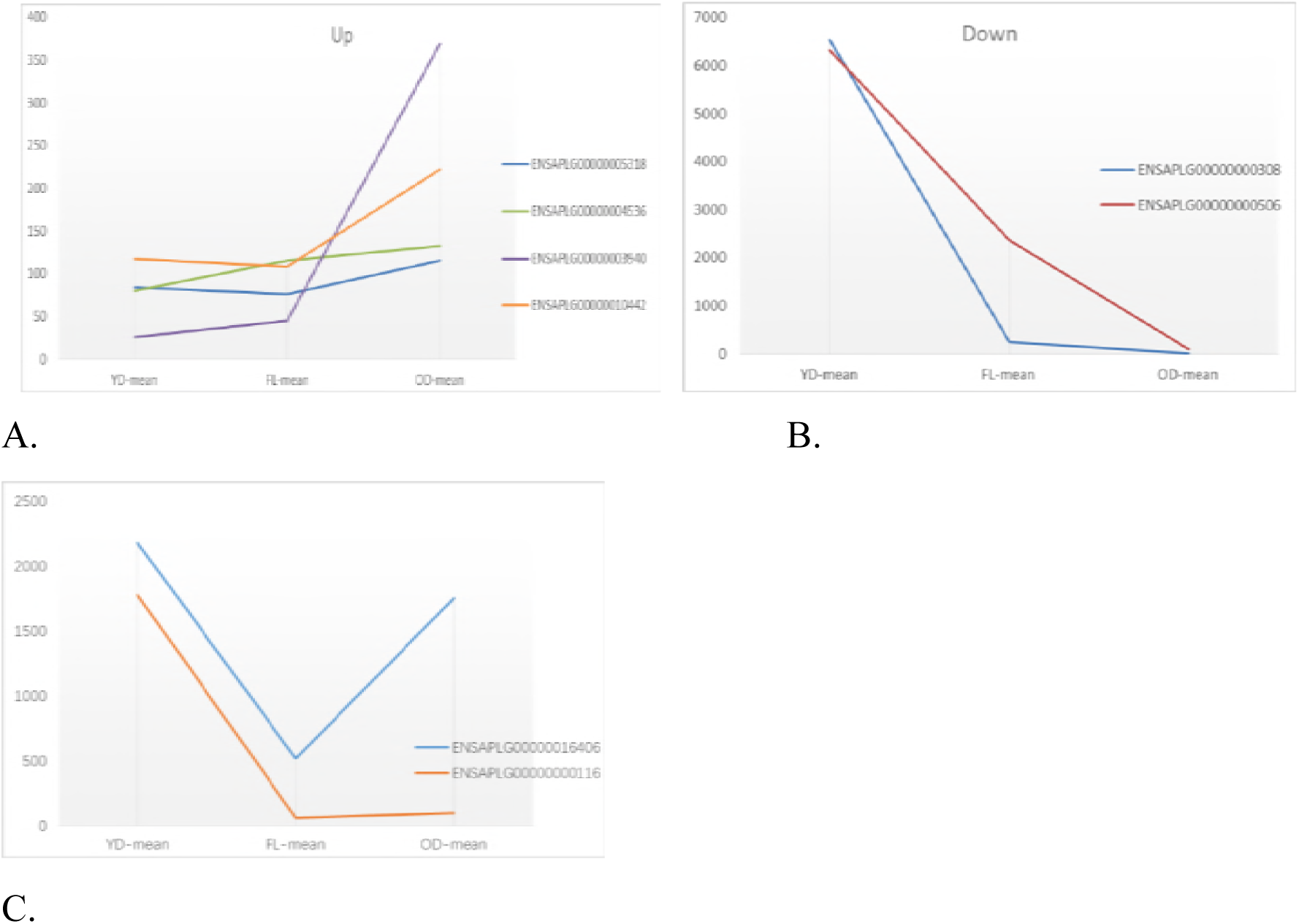
The DEGs exhibit either upregulation, downregulation, swing patterns between life stages. (A) The DEGs with upregulation profile in duck life cycle. (B) The DEGs with downregulation profile in duck life cycle. (C) The DEGs with swing patterns in duck life cycle.

Divergent genes identified in the comparison between OD and FL groups could be clustered into three groups where the ratio of the sum of squares was greater than 80% regardless of the tissue source (Table 2). The most suitable clustering ratio of the sum of squares in young duck tissue sources was 91.1%, and genes within each cluster were significantly downregulated with increasing age. Cluster No. 3 showed the most remarkable downregulation in both transition phases (young to first-laying, and first-laying to old; Fig 1B). Divergent genes from these comparisons showed high fit to the three clusters; with most of their RPKMs in the range 20 to 50, and the divergent genes in clusters with the most genes were expressed at levels 50 to 150 times lower than expression levels of divergent genes in other clusters. The highest fitful clustering from young ducks showed significant downregulation across the three life stages from young ducks to laying ducks, and from laying ducks to old ducks.

**Table 2.** Expression levels in young ducks (YD), first-laying ducks (FL) and old ducks (OD) for genes differentially expressed between FL and OD. “Ratio of SS” indicates the ratio of within-cluster sum of squares to total sum of squares. Different lowercase letters in the same row indicate significant difference at P < 0.05.

| Cluster source | Cluster No. | Number of divergent genes | Ratio of SS | YD | FL | OD |
| --- | --- | --- | --- | --- | --- | --- |
| Young ducks | 1 | 3 | | 2143.02 ± 1124.465 | 2070.87 ± 1174.042 | 446.7 ± 447.237 |
| | 2 | 513 | 91.1% | 40.78<sup>A</sup> ± 89.736 | 40.55<sup>A</sup> ± 104.31 | 24.74<sup>B</sup> ± 87.061 |
| | 3 | 2 | | 6427.87<sup>A</sup> ± 156.381 | 1307.22<sup>AB</sup> ± 1506.741 | 54.15<sup>B</sup> ± 70.125 |
| Laying ducks | 1 | 508 | | 48.37<sup>a</sup> ± 296.671 | 31.58<sup>a</sup> ± 54.516 | 22.1<sup>b</sup> ± 82.223 |
| | 2 | 8 | 81.4% | 743.71<sup>Aba</sup> ± 594.294 | 977.54<sup>Aa</sup> ± 359.135 | 239.49<sup>Bb</sup> ± 191.762 |
| | 3 | 2 | | 4841.29 ± 2087.38 | 2884.43 ± 723.775 | 499.51 ± 559.712 |
| Old ducks | 1 | 2 | | 1791.05 ± 2226.316 | 1896.02 ± 2121.604 | 1188.73 ± 415 |
| | 2 | 501 | 83.4% | 61.77<sup>A</sup> ± 412.346 | 39.82<sup>B</sup> ± 142.274 | 13.78<sup>C</sup> ± 25.909 |
| | 3 | 15 | | 378.42 ± 499.67 | 392.58 ± 447.73 | 323.99 ± 119.69 |

### Function analysis of DEGs

The DEGs from pairwise group were cluster to biological process, cellular component and molecular function (Fig 2). According to the GO enriched results, the number of DEGs showed a significant reduction in the follicular tissues of young ducks, laying ducks and old ducks. Among these DEGs, young ducks have richment genes expression with high level, and the most down-regulation genes appear in transition of YD to FL, and about 300 DEGs were found in cellular process, single organism process, biological regulation and multicellular organism process (Fig 2A). In these listed process, however, only about 10 DEGs were clustered to development process with down-regulation function. A significant difference was found in transition of FL to OD, there are more than 180 DEGs with reduced expression and more than 30 DEGs with up-regulated expression are associated with developmental processes (Fig 2B).

**Fig 2.**
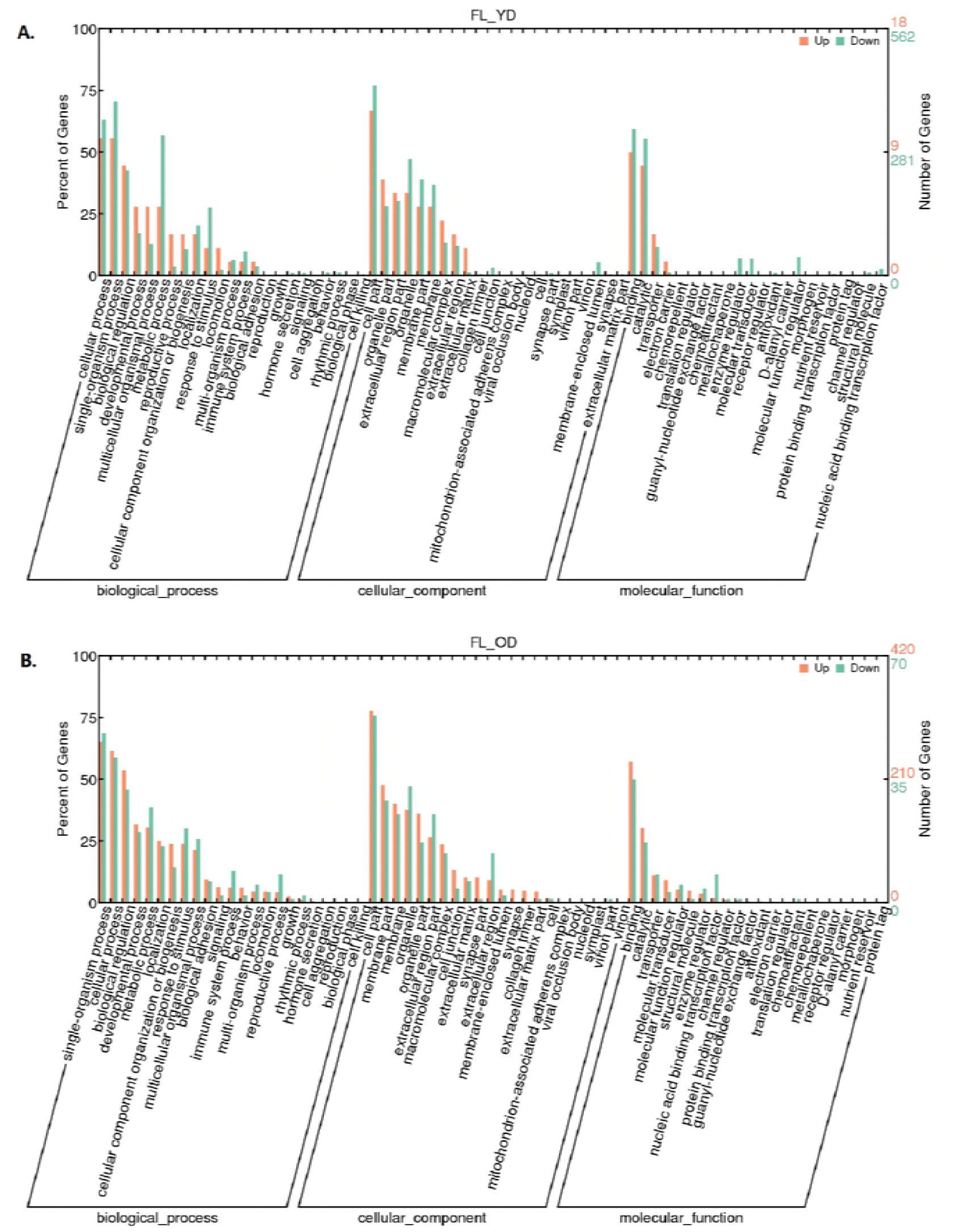
The results of analysis of DEGs by GO in two different transition. (A) The GO analysis result of DEGs from the transition from YD to FL. (B) The GO analysis result of DEGs from the transition from FL to OD.

Further analysis of DEGs by KEGG showed that there was a significant change in the metabolic pathways during the YD to FL transition, and significant changes in the ovarian steroids synthesis pathway and steroid hormone biosynthesis pathway during FL to OD transition (Fig 3). Moreover, the DEGs enriched with the ovarian steroid hormone synthesis pathway and the steroid hormone biosynthesis pathway are consistent with DEGs enriched in the development process of GO enrichment. The synthesis of steroid hormones is closely related to the development of follicles, and also is key biological processes occurring in the theca cells and granulosa cells of the follicle. Based on the results of KEGG, there are seven DEGs from FL comparison with OD group (Fig 4). Among of them, six DEGs showed up-regulation in FL, and the highest one expression level up to 40 times higher than its expression level in OD group. According to the results of KEGG analysis, the only dwon-regulation DEGs is SR-B1 (scavenger receptor B type function as cholesterol transmembrane transfer, and its expression level was reduced by more than twice in the FL group compare to OD group. In the YD group, its expression level is also 35% higher than that of the FL group, but it did not reach statistically significant levels. The only one up-regulation DEG in transition of from YD to FL is EPOX (the member of cytochrome P450 family) play a role of enhancing amino acid metabolism in theca cell of follicle. The other up-regulation DEGs included StAR, CYP17, 3β-HSD, CYP19A1 and CYP1B1 that all of them have higher expression level in YD and FL than in OD.

**Fig 3.**
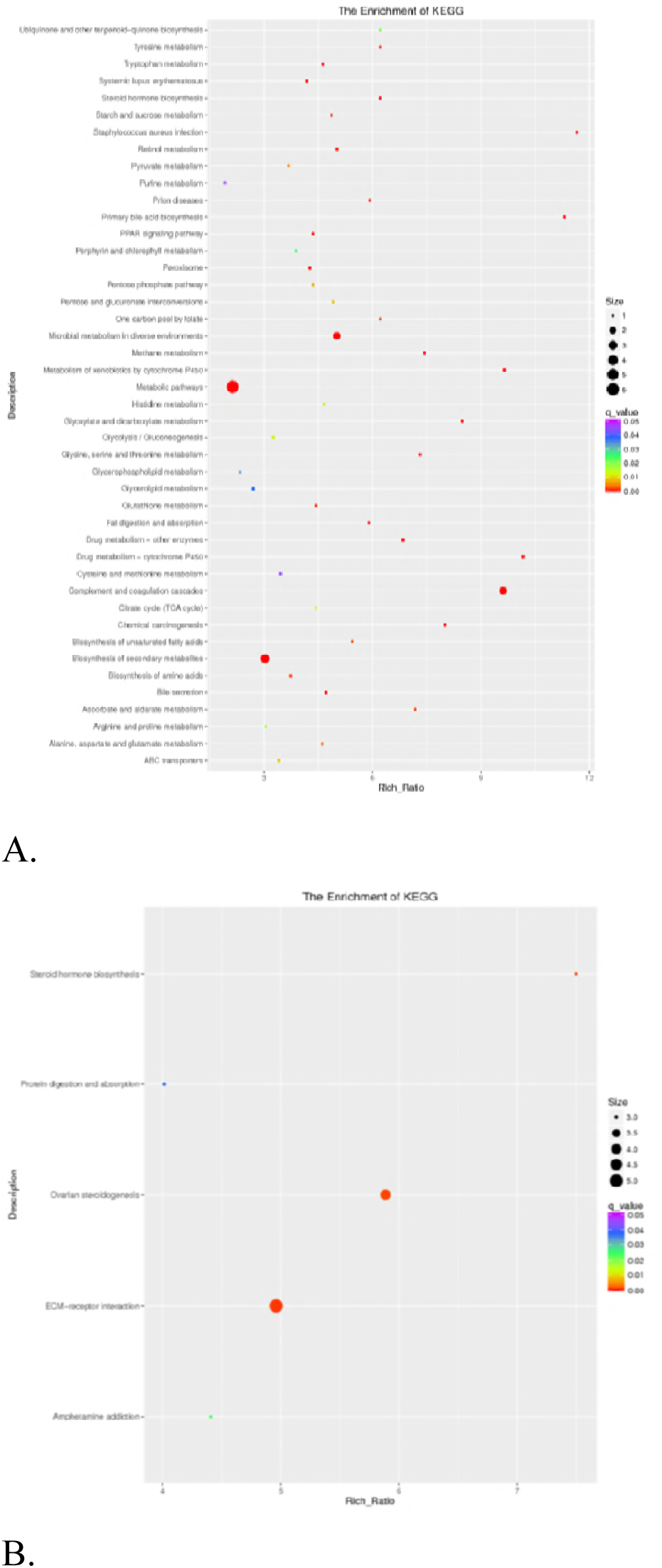
The results of analysis of DEGs by KEGG in two different transition. (A) The KEGG analyzed result of DEGs from the transition from YD to FL. (B) The KEGG analyzed result of DEGs from the transition from FL to OD.

**Fig 4.**
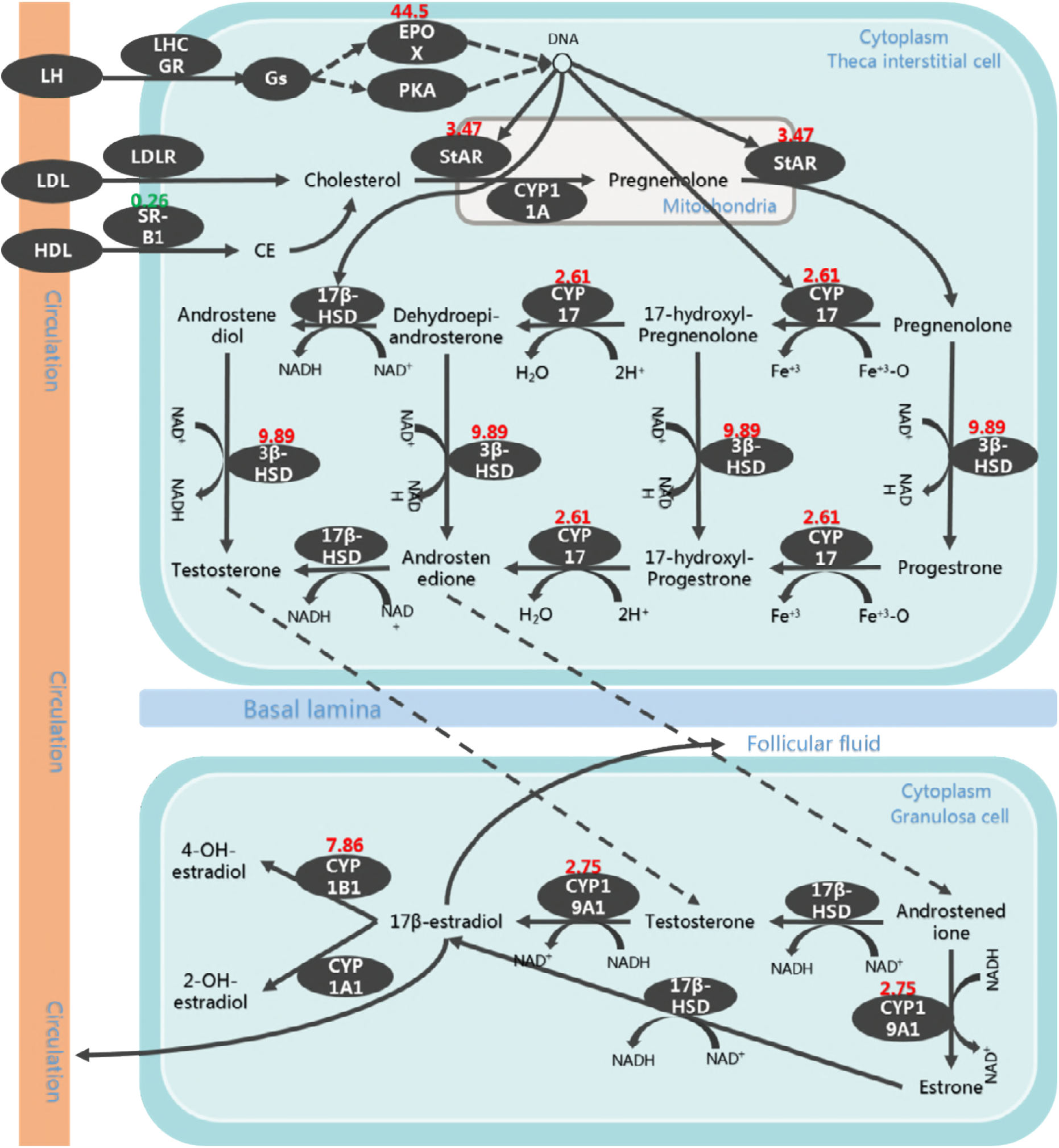
The change of steroid hormone biosynthesis pathway in transition from FL to OD. Note: The red numbers were DEGs’ up-regulated expression folds by FL vs OD. down-regulated DEGs noted with green color.

## DISCUSSION

Some studies have demonstrated that divergent ovarian genes in ducks play a role in reproduction and cluster into different groups based on their expression levels. Ducks comprise the second biggest source of poultry eggs and meat; therefore, development of breeding technologies that improve the quality of duck egg traits is important. Several studies have reported that divergent genes play a role in the development of follicle dominance [20] and ovary development [21] in animals, but no studies have reported the same in birds. Limited knowledge of divergent expression in ducks has prevented the improvement of reproductive efficiency using molecular breeding technology. However, reports on duck and goose genome sequences offer a means of exploring differential gene expression in duck ovaries[22-23]. Using the published duck genome in combination with transcriptome sequencing, we identified 593 coding genes and 14 noncoding genes that differed between young and laying ducks, and 518 coding genes and 93 noncoding genes that differed between laying and old ducks. These results enrich our knowledge of divergent gene expression in duck ovaries and follicle.

Clustering analysis showed that these differential ovarian gene expression levels clustered into three groups with high degrees of suitability. The identified divergent coding genes from the comparison between laying ducks and young ducks were 593. The clustering of these genes’ by their expression levels in young ducks demonstrates a swing distribution, in which most of the genes clustered to Group 2 and a few genes clustered to Groups 1 and 3. Five of the identified genes belonging to Group 1 produce proteins functioning in defense roles, such as Gallinacin-10 and C factor, to improve young duck health, and proteins associated with development, such as Matrix Gla [24], and are only expressed at high levels in young ducks and at no other life stage. The largest clustering group of divergent genes was Group 2. In this group, mean expression levels of divergent genes were broadly similar in different life stages, although the expression levels of many genes coding for functional proteins, such as dehydrogenase and reductase, were affected by time [25]. Therefore, it is predictable that they would be found in the divergent gene pool. The third cluster includes two genes, coding for Hemoglobin subunit alpha-A and RNase, the former of which plays a key role in embryonic transactivation to complete embryonic development in chickens [26]. According to these cluster results, duck ovaries undergoing the transition from the young phase to the laying phase receive addition proteins involved with defense systems; genes related to ovary maturity also show high expression levels during this transition. Although each phase demonstrated different clustering groups, most genes remained in the same groups between phases.

Steroid hormones are related to the development of sex and secondary sexual characteristics of animals, and play an important role in maintaining the reproductive characteristics of birds [27]. The estrogen secreted by the female ovaries is the most important. Previously studies have shown that it has the effect of promoting female follicular development and ovulation [28]. Therefore, their synthesis has been the key research object for researchers to explore the development of animal reproductive organs and reproductive performance. Previously, there have been specific reports on such hormone synthesis pathways [29-30], but in the development of avian follicles, especially in domesticated birds, which genes are regulated by these hormons synthesis have not been reported.

In this study, through DEGs clustering and further functional analysis, we found that most of the genes in the steroid hormone synthesis pathway in duck reproductive organs have been initiated expression from the youth development stage, such as 3β-HSD, StAR, etc., but in reproduction stage specific. Activity of expression in these genes is significant decline after the cessation of follicular maturation has not been reported. In addition, this study found that the EPOX protein in the ovarian steroid hormone synthesis pathway has a specific high expression in the reproductive stage during the entire reproductive life of the duck. EPOX is a member of the cytochrome P450 superfamily of enzymes [31]. The enzymes are oxygenases which catalyze many reactions involved in the synthesis of cholesterol, steroids and other lipids. Its high expression in the laying stage of ducks can provide a large amount of raw materials for the synthesis of reproductive hormones, ensuring the normal development of follicular development activities. Ovarian follicles mainly increase and maintain development in the youth stage [32], and follicular development and maturation activities begin in the laying stage, indicating that EPOX is a key gene regulating duck follicular development, and its expression can increase with follicular development in theca and granulosa cells of birds follicle. Moerover, EPOX have the ability to enhance amino acid metabolism regulates DNA, and indirectly promoting the expression of other key genes in the steroidogenic synthesis pathway [33], such as StAR, CYP17 and 3β-HSD, to provide sufficient organic molecules for the synthesis of estrogen required for duck follicular development. The results of this study also showed that steroidogenic synthesis pathway have same key DEGs in duck, but the expression mode of key DEGs are different, such as when the expression level of EPOX decreased from 100% of the reproductive stage to 2%, the expression of indirectly regulated gene was also significantly reduced. However, EPOX did not have a significant effect on the regulation of StAR, CYP17 and 3β-HSD expression levels before laying eggs.

In conclusion, we clustered differentially expressed functional genes and their function analysis, and found that DEGs could be categorized into three groups. The main functional class of DEGs associated with follicular development is the steroid hormone synthesis pathway. Further analysis indicated that StAR, CYP17, EPOX, 3β-HSD, CYP1B1 and CYP19A1 in the steroid synthesis pathway are key factor for maintenance of follicular maturation and reproductive activity, among which EPOX is a key gene for time special highly expression during egg laying stage, and other key regulatory genes’ highly expression showed in young and laying stage and lower expression showing with follicular development stopping.

## ACKNOWLEDGMENTS

We thanks the Zhejiang Zhuowang Agriculture Sci-Tech Limited Co,. offering samples needed in this study and the Annoroad Research Team providing data analysis support. This study was supported by the Earmarked Fund for National Waterfowl-industry Technology Research System (number CARS-43-2 and CARS-42-2) and the Zhejiang Major Scientific and Technological Project of Agricultural (livestock’s) Breeding (2016C02054-13).

